# Reduction in activity and abundance of mitochondrial electron transport chain supercomplexes in pulmonary hypertension-induced right ventricular dysfunction

**DOI:** 10.1101/2024.03.08.584016

**Authors:** Wenzhuo Ma, Peng Zhang, Alexander Vang, Alexsandra Zimmer, Scarlett Huck, Preston Nicely, Eric Wang, Thomas J Mancini, Joseph Owusu-Sarfo, Clarissa F. Cavarsan, Andriy E. Belyvech, Kenneth S. Campbell, Dmitry Terentyev, Gaurav Choudhary, Richard T. Clements

## Abstract

Pulmonary hypertension (PH) results in RV hypertrophy, fibrosis and dysfunction resulting in RV failure which is associated with impaired RV metabolism and mitochondrial respiration. Mitochondrial supercomplexes (mSC) are assemblies of multiple electron transport chain (ETC) complexes that consist of physically associated complex I, III and IV that may enhance respiration and lower ROS generation. The goal of this study was to determine if mSCs are reduced in RV dysfunction associated with PH. We induced PH in Sprague-Dawley rats by Sugen/Hypoxia (3 weeks) followed by normoxia (4 weeks). Control and PH rats were subjected to echocardiography, blue and clear native-PAGE to assess mSC abundance and activity, and cardiomyocyte isolation to assess mitochondrial reactive oxygen species (ROS). mSC formation was also assessed in explanted human hearts with and without RV dysfunction. RV activity of CI and CIV and abundance of CI, CIII and CIV in mitochondrial mSCs was severely reduced in PH rats compared to control. There were no differences in total CI or CIV activity or abundance in smaller ETC assemblies. There were no changes in both RV and LV of expression of representative ETC complex subunits. PAT, TAPSE and RV Wall thickness significantly correlated with CIV and CI activity in mSC, but not total CI and CIV activity in the RV. Consistent with reduced mSC activity, isolated PH RV myocytes had increased mitochondrial ROS generation compared to control. Reduced mSC activity was also demonstrated in explanted human RV tissue from patients undergoing cardiac transplant with RV dysfunction. The right atrial pressure/pulmonary capillary wedge pressure ratio (RAP/PCWP, an indicator of RV dysfunction) negatively correlated with RV mSC activity level. In conclusion, reduced assembly and activity of mitochondrial mSC is correlated with RV dysfunction in PH rats and humans with RV dysfunction.

## Introduction

Increased afterload associated with pulmonary hypertension (PH) results in right ventricular dysfunction associated with numerous right ventricular (RV) mitochondrial and metabolic changes[39, 41]. Altered RV mitochondrial function and metabolism occurs in multiple animal models of PH and humans [41]. These include alterations in mitochondrial substrate utilization, content and architecture changes, respiration, and increased reactive oxygen species (ROS) generation which are all known to affect cardiomyocyte contractile function (rev in [34, 41]). In response to persistent pressure overload, the RV undergoes metabolic reprogramming with reduced oxygen consumption, respiration, and electron transport chain (ETC) activity as shown in animal models of PH and RV dysfunction [8, 13, 17, 40]. There are also global metabolic shifts to glycolysis in both animal models of PH and patients [18, 27, 34] as well as increases in glutaminolysis[28] , both likely to maintain energy reserves in the context of reduced oxidative phosphorylation. In addition, there is evidence suggesting ROS is elevated in RV cardiomyocytes presumably from mitochondrial sources which likely contributes to pathology associated with right heart failure in PH [31, 35, 43]. RV dysfunction in PH eventually results in RV failure and death in PH patients, however there are no therapies targeting RV dysfunction or metabolic alterations, and no therapies for left heart failure have shown efficacy for treatment of RV failure [29]. Therefore, it is urgent to uncover novel pathological mechanisms that may provide new therapeutic targets for RV maladaptation and failure in PH.

The ETC produces the proton gradient across the mitochondrial inner membrane necessary for ATP production and is also responsible for mitochondrial ROS generation. It is increasingly recognized that ETC complex structures are dynamic and different assemblies of the ETC can impact respiration and ROS generation [1, 20]. Mitochondrial ETC supercomplexes (mSC) are composed of individual complexes of the electron transport chain that physically associate into supercomplexes consisting of assemblies of complex I, III and IV[20]. There is increasing evidence that these assemblies promote more efficient respiration via increased respiration while potentially reducing ROS generation through less electron slippage and more efficient transfer between the complexes[1].

We hypothesized that there are reduced mSC assemblies in the RV associated with RV dysfunction in PH. In this paper, we report that mSC activity and assembly are decreased in the RV in the sugen/hypoxia (SuHx) model of pulmonary arterial hypertension. Activity of mSC, but not activity of smaller or individual assemblies, correlated with severity of RV dysfunction and hypertrophy in PH. In addition, we found that RV mSC activity was also reduced in human hearts with RV dysfunction related to group II PH.

### Materials and Methods Animals

Animal use was approved by the Institutional Animal Care and Use Committee of the Providence VA Medical Center in compliance with the National Institutes of Health guideline for the care and use of laboratory animals.

### Human heart tissues and patient characteristics

RV myocardium from 24 subjects (12 male,12 female) was obtained from the Gill Cardiovascular repository at the University of Kentucky. 5 patients were excluded from analysis, 2 due to diagnosis of ischemic heart failure and 3 due to right heart catheterization (RHC) data either lacking or recorded >1 year before transplant. All patients had left heart failure, and most had elevated pulmonary artery pressures consistent with group II PH. All procedures were approved by the University of Kentucky IRB and the samples collected and stored at time of transplant as previously described [2].

### SU5416/Hypoxia rat model of PH

Adult male (12) and female (12) Sprague-Dawley rats (175-200 g) were purchased from Charles River (Wilmington, MA). Animals were divided into 2 groups: normoxia (Con; n=12) and PH (n=12). PH was induced by a single subcutaneous injection of SU5146 (Cayman Chemical) 20 mg/kg and placed into a normobaric hypoxia chamber (10% FiO2; Biospherix, Ltd, Parish, NY) for 3 weeks [37]. Con animals were injected with diluent subcutaneously and exposed to normoxia for the entirety of the study (7 weeks). After 3 weeks hypoxia, the PH group was returned to normoxia for the remaining 4 weeks similar to our previous studies [6, 38]. 2 PH animals died early during the course of the study prior to sacrifice, and were not included in any analyses. An additional PH animal was excluded from correlation analysis due to lacking echocardiographic data caused by arrest during anesthesia for final echocardiography, but the heart was rapidly excised and tissue was included in molecular studies (mSC assembly and activity). One control animal (F) was excluded from all analyses as there was an exceptionally low mitochondrial protein yield (<40% vs nearest control or PH samples) likely related to tissue sampling or technical error and was removed from all analyses following positive outlier detection using Dixon’s Q-test.

### Echocardiography

Echocardiography was performed at 7 weeks similar to our previous publications [6]. Subsequently, rats were anesthetized with isoflurane, and RV and LV tissue were harvested, weighed and snap frozen for subsequent molecular analysis.

### Mitochondrial isolation and BN/CN-PAGE

Approximately 50 mg of frozen RV and left ventricular (LV) myocardium was minced on dry ice and placed in 1mL ice-cold mitochondria isolation buffer (MIB) (0.28M sucrose, 10 mM HEPES and 2 mM EDTA, PH 7.4) supplemented with protease and phosphatase inhibitors (Sigma). Tissue was homogenized in MIB with a Dounce homogenizer, using 15 strokes each of loose and tight pestles. Mitochondria were isolated by differential centrifugation according to standard methodology. An initial 2,000g x 5min spin to isolate nuclei, followed by a 13,000g x 10 min spin to isolate mitochondria. Mitochondrial pellets were resuspended in 1 ml MIB and were washed twice with spins at 13,000g. Mitochondrial pellets were suspended in 100 μL MIB and subjected to BCA protein assay to assess total mitochondrial mass.

Mitochondrial pellets were then solubilized in 1% digitonin in MIB buffer (1μL/μg) and subjected to a second protein BCA assay (Thermo). Equal amounts of mitochondrial protein were resuspended in Native-PAGE sample buffer (Invitrogen) supplemented with 1% Digitonin (Invitrogen) and 0.25% Coomassie G250. Equal μg of mitochondria (∼10 μg) were loaded on Invitrogen 4-16% Native-PAGE gels. Gels were run for 60 min at 150V with Invitrogen Native-PAGE anode and dark blue cathode buffers followed by 250V with light blue cathode buffer (Invitrogen) for 180 min according to manufacturer’s instructions. Gels were blotted to nitrocellulose membranes and probed for complex I (Ndufa9; 1:1000 Abcam), complex IV (Cox IV, 1:1000; Abcam) and complex III (UQCRFS1; 1:1000 Abcam), in that order with stripping in between using 10% SDS/ 2% 2-mercaptoethanol stripping buffer at 37**°**C.

One-dimensional electrophoresis in SDS-PAGE was performed using whole cell lysates collected following initial dounce homogenization in MIB, protein normalization with BCA and solubilization with 4X SDS sample buffer. 0.5 mg of whole tissue lysates were separated using SDS-PAGE and transferred to nitrocellulose membranes that were processed by immunodetection, using OXPHOS cocktail (Abcam, 1:1000) followed by strip and reprobe for HSP60 (Cell signaling technology,1:1000).

### Clear-Native PAGE and mitochondrial respiratory chain supercomplexes activity assessment

Clear-Native PAGE was performed similarly to Blue-Native PAGE with modifications as in [16]. Gels were run for 60 min at 150V with light blue cathode buffer followed by 250V with clear cathode/running buffer for 180 min. The gel was removed and placed in ice cold water followed by incubation with complex IV buffer (0.5 mg/mL Diaminobenzidine, 1 mg/mL cytochrome c in phosphate buffer (38 mM Na_2_HPO_4_ and 12.3 mM NaH_2_PO_4_, pH 7.4)) or complex I buffer (0 .1 mg/mL NADH, 2.5 mg/mL nitrotetrazoleum blue chloride in 2 mM Tris, pH 7.4). Gels were incubated in 20 mL of their respective solution, monitored for color development 30-60 min, and the reaction stopped with 10% acetic acid, and washed in dH_2_O.

### Cardiomyocytes isolation from control and PH rats

Right ventricular myocytes were isolated from control and PH rats as described previously with minor modification[22, 38]. Sprague-Dawley rats of control and PH were euthanized with isoflurane. Hearts were immediately excised, trimmed, and retrogradely perfused at 8 mL/min for 2 min with Ca^2+^-free Krebs-Henseleit Buffer (KHB) (118 mM NaCl, 4.7 mM KCl, 1.2 mM MgSO_4_, 1.2 mM KH_2_PO_4_, 25 mM NaHCO_3_, 11 mM Dextrose, 8.4 mM HEPES) at 37°C, perfused with enzyme buffer (0.3 mg/mL collagenase II, 0.3 mg/mL hyaluronidase type II, and 50 μM CaCl_2_ in KHB) for 15-20 minutes. After perfusion, LV and RV were separated and RV tissue was minced and placed in enzyme buffer (0.6 mg/mL trypsin IX, 0.6 mg/mL deoxyribonuclease, and 500 μM CaCl_2_ in KHB) in a 50 mL centrifuge tube for further digestion at 37°C for 15 min in a shaking water bath. After further digestion, RV cells suspension were filtered with 200 μm mesh into a 10 mL washing medium (KHB, DMEM/PS, 1:1 ratio) centrifuge tube, then centrifuged at 20g for 3 minutes. The pellet was resuspended with another 10 mL washing buffer and RV cells were precipitated with 0.6% BSA for cardiomyocyte purification. Then RV cells were plated on laminin-coated (Corning) dishes and after 2 hours the medium was changed to culture medium (2 g/L albumin, 2 mM L-carnitine, 5 mM creatine, 5 mM taurine,0.1 μM insulin, and 1% P/S in Medium199).

### Cardiomyocytes mitochondrial ROS and membrane potential measurement

Cardiomyocytes were infected with adenovirus encoding the matrix-targeted ROS biosensor matrix-HyPer7 as described by Pak et al. [26]. Following 24 hours infection cells were placed in Tyrode’s solution (140 mM NaCl, 5.4 mM KCl, 0.53 mM MgCl_2_, 0.33 mM NaH_2_PO_4_, 5.5 mM Glucose, 25 mM HEPES, 10 mM Na Pyruvate, 1.2 mM CaCl_2_.) and imaged on a Zeiss LSM 800 confocal microscope. The probe is ratiometric and cells were sequentially imaged with 400 nm excitation/535 nm emission and 488 nm excitation/535 nm emission every 30 s. After 2-4 min baseline imaging, cells were treated with 5 mM dithiothreitol (DTT) for 3-4 minutes to establish minimum fluorescence ratio and then 6 mM H_2_O_2_ similar to our previous publications [5]. % oxidation was calculated as the ratio of 488/400 excitation normalized to minima (DTT) and maxima (H_2_O_2_).

### Statistical Analysis

Statistical tests were performed with microsoft excel (unpaired t-tests or pearson’s correlation as indicated) or with R for myocyte ROS studies using two-level random intercept model with Tukey’s posthoc as previously described [5, 36] . All data is presented as mean ± SEM unless otherwise indicated. Data were subjected to unpaired two-tailed t test or Pearson’s Correlation as indicated. P<0.05 was considered significant.

## Results

### Sugen5416/hypoxia-induces pulmonary hypertension and RV dysfunction in rats

PH was induced in male and female Sprague-Dawley Rats (Charles River) via Sugen/5416 injection and 3 week hypoxia treatment (10% FiO_2_) followed by standard housing (4 weeks) as in our previous studies[6, 25]. Control animals received placebo injection, and normoxic housing (n=11) (**Figure 1A**). Induction of PH in rats was verified by transthoracic echocardiogram and heart weights at 7 weeks by reduced pulmonary acceleration time (PAT; Con 27.90 ±1.40 ms vs PH 17.23 ± 0.53 ms ) and RV dysfunction as assessed by tricuspid annular plane excursion (TAPSE;Con 2.0± .08 mm vs PH 1.11±.10 mm) , and cardiac output (CO) (Con 93.67 ± 5.62 mL/min vs PH 58.93 ± 5.48 mL/min, and increased RV hypertrophy as shown by Fulton Index (RV/LV+S) and RV thickness (**Figure 1 B,C**).

**Figure 1:**
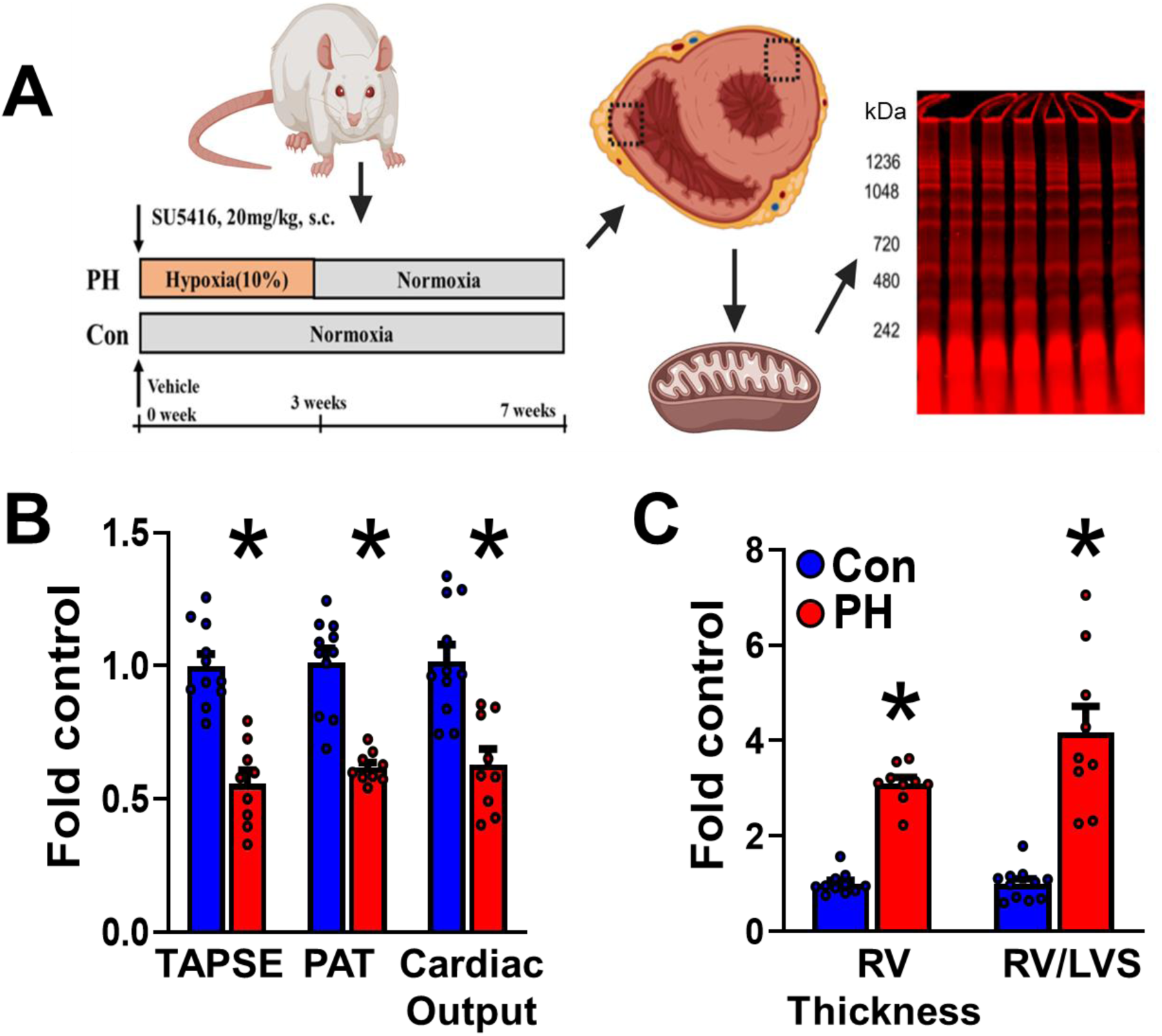
RV dysfunction and hypertrophy in Sugen/hypoxia rats. **A)** Schematic of experimental design: Rats subjected to Sugen/Hypoxia (PH) or normoxia (Con) for 3 weeks followed by 4 weeks normoxia, echocardiography, mitochondrial isolation from RV and LV and Blue or Clear-Native PAGE and isolation RV myocytes. **B,C**) Echocardiographic analysis at sacrifice demonstrates RV dysfunction and hypertrophy as shown by (**B**) TAPSE, PAT, and cardiac output as well as (**C**) Fulton Index and RV wall thickness in PH rats. Normalized to control. *p<.05 as indicated, N=11 Control and 9 PH.

### PH reduces mitochondrial supercomplex activity in the right ventricle

Isolated mitochondrial protein from the RV and LV of control and PH rats were subjected to CN-PAGE which separates large native molecular weight complexes from ∼200 kDa to >2 mDa. Complex I and IV specific activity of intact ETC complexes was visualized via activity-dependent colorimetric modification of substrate (complex I, CI: Nitrotetrazolium Blue Chloride, violet; and complex IV, CIV: Diaminobenzedene, brown). Protein load was assessed via far-red licor imaging of Coomassie stain (**Figure 2A,B**). Supercomplexes were identified as overlap of CI and CIV in the region > ∼1000 kDa based on expected molecular weight of CI/CIII_2_/CIV_1-n_ assemblies. **Figure 2C** shows a higher contrast black and white image of the gels in 2A clearly demonstrating decreases in mSC assemblies in the RV. Quantitation of the mSC region normalized to total mitochondrial protein load demonstrated significant decreases in RV mSC associated CI and CIV activity in PH rats. There were also decreases in CI and CIV activity in the mSC region in LV mitochondria, but to a lesser extent than alterations in RV (**Figure 2D**). There were no changes in total CI and CIV activities (sum of mSC and smaller ETC assemblies visualized in lane) in the RV or LV (**Figure 2E**).

**Figure 2:**
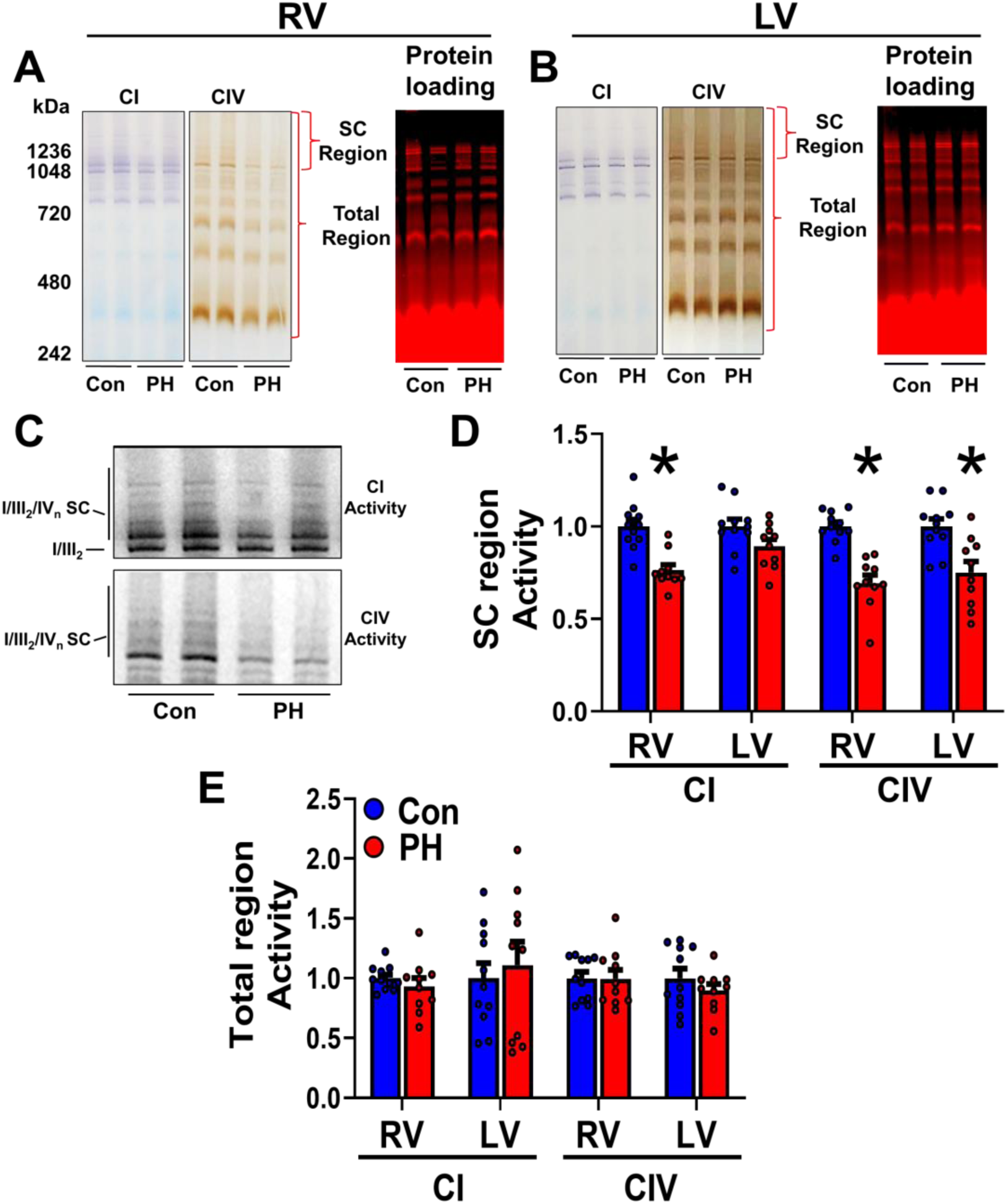
Activity of mSC is reduced in PH rats. Representative CN-PAGE of rat **(A)** RV and (**B**) LV (right) mitochondria for CI (purple) and CIV (brown) activity in isolated mitochondria. Total protein load of mitochondrial lysates is shown by licor imaging of Coomassie stain (red) Activity in the mSC region is lower in PH rats. **C**) Zoomed inset region of SC shown in A in black and white to enhance contrast. **D,E**) Quantitation of gels in A show CI and CIV activity significantly decreased in the mSC region compared to control in the RV and CIV mSC activity decreased in the LV. Total activity (entire activity signal in lane) was unchanged. *p<.05 as indicated, N=11 control and 10 PH.

### Supercomplex activity negatively correlates with severity of RV dysfunction in PH

Tricuspid annular plane systolic excursion (TAPSE) which is lower in PH, was positively correlated with both CI (**Figure 3A**) and CIV (**Figure 3B**) activity in the mSC region. RV wall thickness (**Figure 3 C,D**) negatively correlated with CI and CIV mSC activity. CI and CIV activity in the mSC region only also correlated with derangements in PAT, cardiac output and Tricuspid E/E’ (**Table I**), indicating reduced mSC with increased severity of PH-induced RV systolic and diastolic dysfunction. There was no correlation between total CI and CIV activity with any of the RV structural and functional parameters (**Table I**).

**Figure 3:**
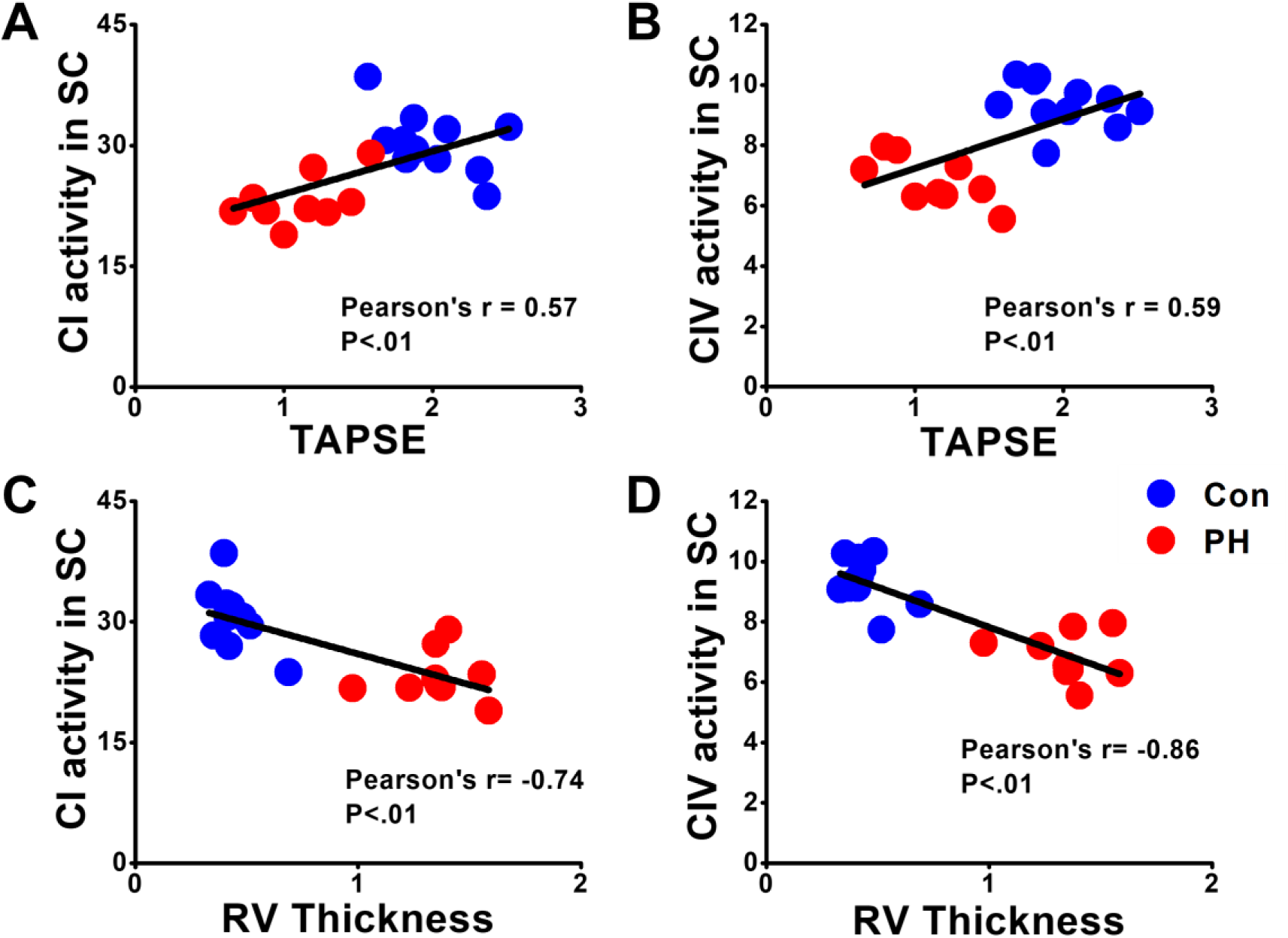
Correlation of RV mSC activity with indices of PH severity in SuHx rats. (**A**) CI and (**B**) CIV activity in mSC positively correlates with RV function as measured by TAPSE. **C,D**) CI (**C**) and CIV (**D**) activity in mSC negatively correlates with RV wall thickness. N=11 Control and 9 PH.

**Table 1:** Pearson correlation of SC region and total CI and CIV activity with hemodynamic indicators of PH severity including PAT, TAPSE, RV thickness, CO and tricuspid E/E’. Significance determined by Pearson’s correlation, N = 11 control and 9 PH.

| RV<br>Parameters | SC<br>CI activity |  | Total<br>CI activity |  | SC<br>CIV activity |  | Total<br>CIV activity |  |
| --- | --- | --- | --- | --- | --- | --- | --- | --- |
|  | r | P | r | P | r | P | r | P |
| PAT | 0.68 | <0.01 | 0.22 | 0.36 | 0.69 | <0.01 | -0.10 | 0.67 |
| TAPSE | 0.57 | <0.01 | 0.30 | 0.19 | 0.59 | <0.01 | 0.19 | 0.43 |
| RV<br>Thickness | -0.74 | <0.01 | -0.34 | 0.14 | -0.86 | <0.01 | -0.02 | 0.93 |
| Cardiac<br>Output | 0.55 | <0.05 | -0.25 | 0.28 | 0.60 | <0.01 | 0.17 | 0.47 |
| Tricuspid<br>valve E/E' | -0.63 | <0.01 | -0.08 | 0.7 | -0.60 | <0.01 | -0.004 | 1.0 |

### Supercomplex assembly is decreased in PH rats

Assembly of RV and LV myocardial ETC mSCs was assessed using blue native-PAGE and immunoblot with the CI, CIII and CIV constituent proteins NDUFA9, UQCRFS1 and COXIV. Representative blots of CI, CIII, and CIV assemblies from RV and LV are shown in **Figure 4A**. As shown in **Figure 4B**, RV mitochondrial mSCs consisting of CI (NDUFA9), CIII (UQCRFS1) and CIV (COXIV) were significantly decreased in PH rats compared to controls (**Figure 4A** boxed regions labeled I/III_2_/IV_n_ SC), consistent with reduced CI and CIV activity in mSC. There were also decreases in CIII and CIV in mSC in the LV but to a lesser extent than RV. There were no changes in total assembled ETC complexes inclusive of smaller non-SC assemblies in the RV mitochondria, however total assembly of CIII was reduced in LV mitochondria. (**Figure 4C**).

**Figure 4:**
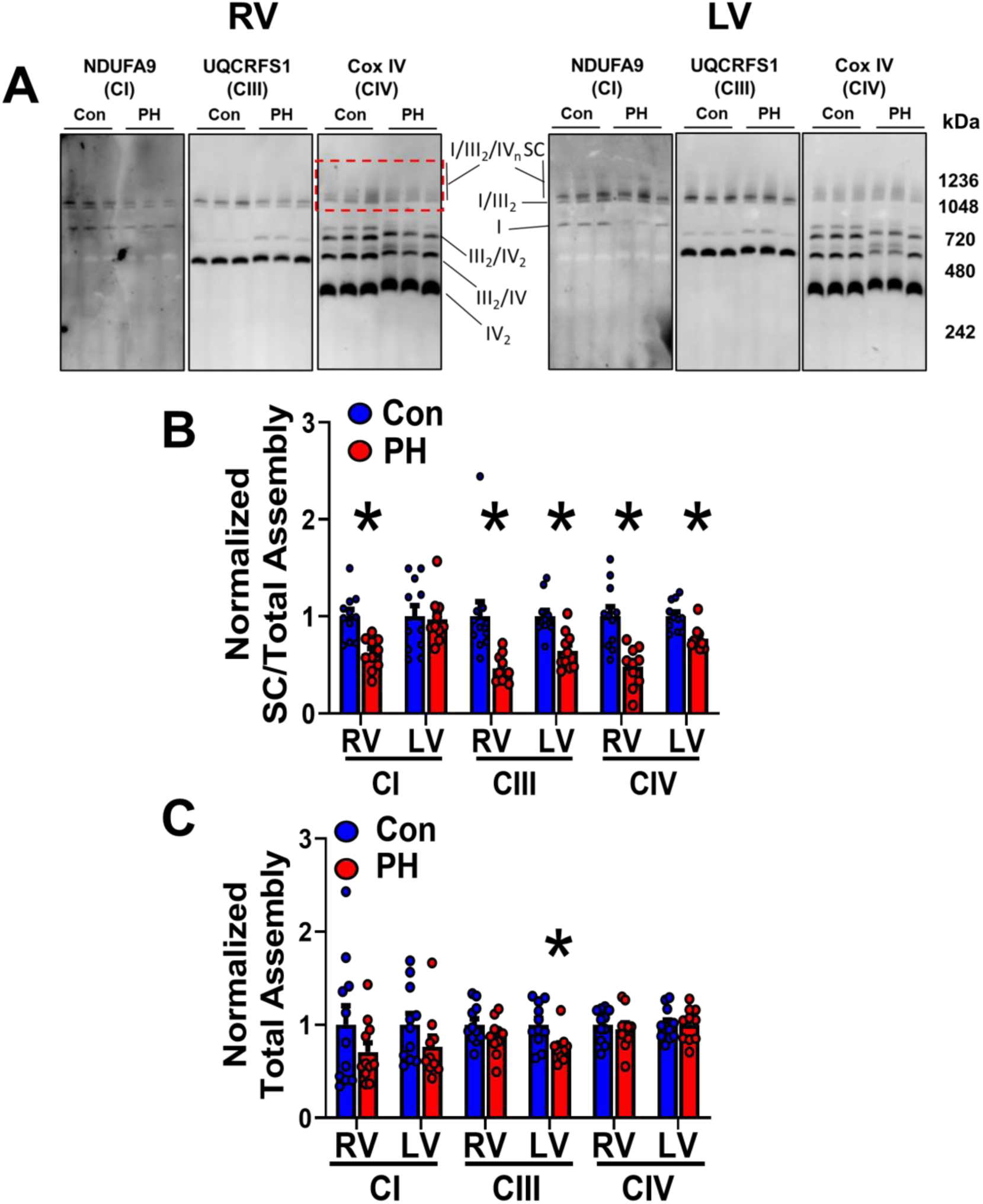
Assembly of mSC is reduced in RV and LV of PH rats. **A**) Representative immunoblots of isolated RV and LV mitochondria subjected to BN-PAGE and immunoblot for NDUFA9 (CI), UQCRFS1(CIII), and CoxIV (CIV). Red dotted box indicates SC region of blot for quantitation. Designations of ETC assemblies and multimers (based on weight and comigration) denoted as complex number and subscript. **B,C**) Quantitation of BN-PAGE density in mSC (boxed region) and total signal (entire lane) of blots shown in A. mSC assembly was greatly reduced in the RV and to a lesser extent in LV in PH animals with minimal changes in total complex . *p<.05 t-test. N=11 control and 10 PH.

### Expression of mitochondrial electron transport chain protein subunits are unchanged in RV and LV myocardium

Mitochondrial ETC protein expression was assessed via SDS-PAGE of total protein lysates with an antibody cocktail recognizing representative ETC complex subunits: NDUFB8 (CI), SDHB (CII), UQCRC2 CIII, MTCO1 (CIV) as well as ATP5A (CV) (**Figure 5A**). There were no changes of ETC protein expression in PH RV and LV compared with control when normalized to the mitochondrial matrix localized HSP60, indicating no selective decreases in ETC proteins in PH (**Figure 5B,C**). Additionally, when expression levels were normalized to cardiac troponin I, results were similar to HSP60 normalization indicating that mitochondrial protein content was similar in control and PH animals (data not shown).

**Figure 5:**
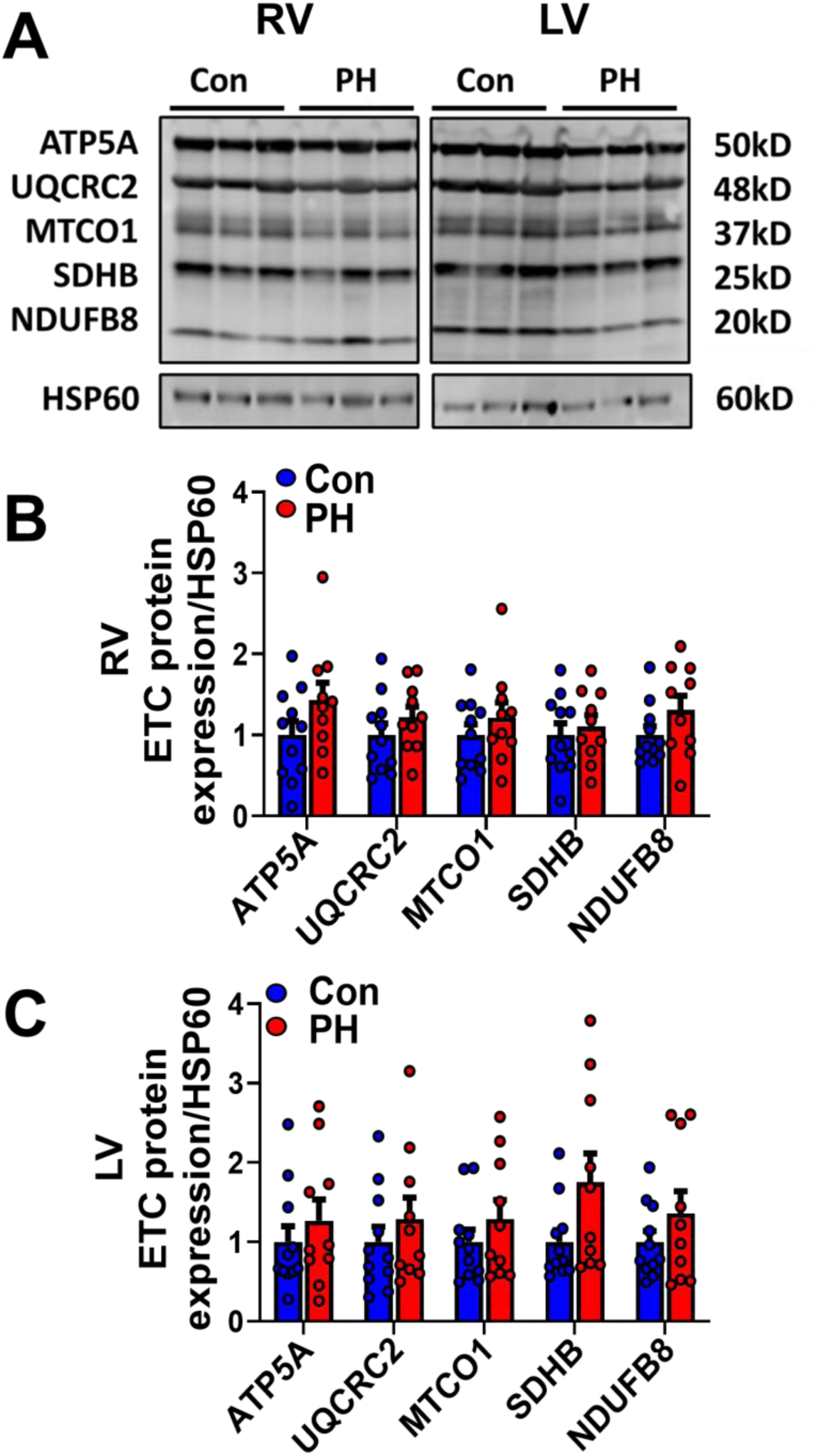
Individual ETC protein subunit expression is not reduced in PH. **A**)Representative SDS-PAGE and immunoblot of whole tissue RV and LV lysates probed for individual representative subunits of ETC complexes (OXPHOS antibody cocktail) blot of mitochondrial ETC proteins. **B,C**) Quantitation of blots in A show no expression changes in representative ETC proteins in mitochondria when normalized to HSP60 in both RV and LV. *p<.05 t-test. Indicated. N=11 control and 10 PH.

### PH increases mitochondrial ROS in RV myocytes

Isolated RV myocytes from control and PH rats were infected with Adv-matrix targeted HyPer7, a ratiometric GFP-based ROS sensor [26]. A high resolution image of a matrix HyPer7 expressing RV cardiomyocyte is shown in **Figure 6A** demonstrating specific mitochondrial localization when excited with 488 nm (green panel) as well as 400nm (pseudo-colored red). **Figure 6B** shows merged green/red images of 488/400 matrix-HyPer7 signal in representative RV cardiomyocytes imaged with lower resolution during experiments showing baseline, DTT, and H_2_O_2_ treatment. The ratio of emission intensity at 535 nm with 488:400 excitation increases (more green) when oxidized (H_2_O_2_) and decreases (more red) when reduced (DTT). PH significantly increases ROS in isolated RV myocytes from SuHx rats as shown in a representative experiment with the 488/400 ratio normalized to fully reduced (DTT:0%) and fully oxidized (H_2_O_2_:100%) treatment is shown in **Figure 6C**. Quantitation of multiple experiments demonstrate RV PH myocytes have significantly increased ROS vs control cells (**Figure 6D**).

**Figure 6:**
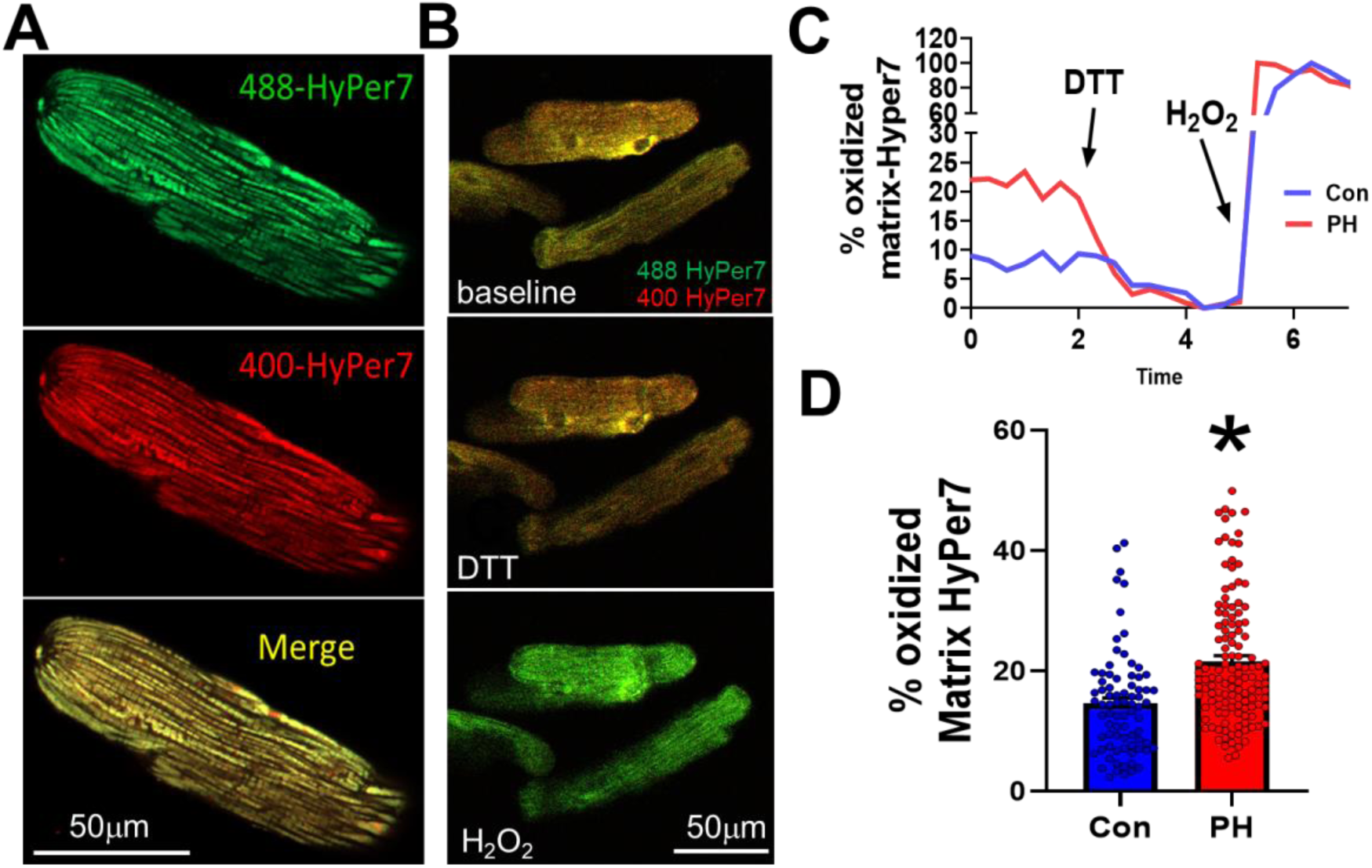
RV myocytes isolated from SuHx PH rats have elevated mitochondrial ROS. **A**) Representative hig resolution image of mitochondrial matrix-HyPer7 expression in an RV cardiomyocyte. Matrix-HyPer7 localized to mitochondria: 488 excitation (green), 400 excitation (pseudo-colored red), and merged signal (yellow). **B**) Low resolution representative images from an experiment showing merged 488ex (green) and 400ex (red) at baseline and following DTT and H_2_O_2_ treatment. **C**) Representative time course of matrix-HyPer7 experiment with normalization to fully reduced ratiometric signal (DTT:0%, min) and fully oxidized signal (H_2_O_2_:100%, max) in control and PH RV cardiomyocytes. **D**) Quantitation of multiple experiments shows PH significantly increased ROS as measured by matrix-Hyper7. * p< .05 as using two-level random intercept model with Tukey’s posthoc test as described in methods. n=78 control and 126 PH RV cardiomyocytes from 6 control and 6 PH rat isolations. *P<.05

### Mitochondrial supercomplex activity negatively correlates with indices of RV dysfunction in explanted human hearts

Patient demographic and RV functional data from right heart catheterization are provided in **Table 2**. Patients diagnosed with ischemic cardiomyopathy were excluded due to potential ischemic effects on the RV independent of increased afterload due to PH. All patients had severely impaired LV function (average EF = 22.4±1.6%, n=16) and most displayed significantly higher mean pulmonary artery pressure (mPAP = 30 mmHg) and pulmonary capillary wedge pressure indicative of Group II PH. The right atrial pressure to pulmonary capillary wedge pressure (RAP/PCWP) ratio has been used as a robust clinical indicator of RV dysfunction in PH associated with transition from compensated to decompensated RV function[7, 14]. As RV dysfunction progresses into failure it cannot maintain cardiac output and RAP increases, and thus RAP/PCWP ratio. A representative CN-PAGE activity analysis is shown in **Figure 7A** showing reduced CI and CIV activity in mSC’s of patients with RV dysfunction (high vs low RAP/PCWP). Correlation analysis with CI and CIV activity in mSC negatively correlated with RAP/PCWP ratio consistent with reduced mSCs in RV dysfunction (**Figure 7B,C**). There was no correlation with mPAP (data not shown) as mPAP is subject to additional influences including significant compensation that has not progressed to overt RV dysfunction (data not shown). There was no correlation with total CI and CIV activity inclusive of smaller CI and CIV assemblies (**Figure 7D,E**). Finally, there was no correlation with overall mitochondrial ETC subunit protein expression with RV function as assessed by western blot of individual ETC subunit proteins (**Figure 7F** and data not shown). Therefore, humans with RV dysfunction display similar derangements in mSC activity as the rat PAH Sugen/hypoxia model.

**Figure 7:**
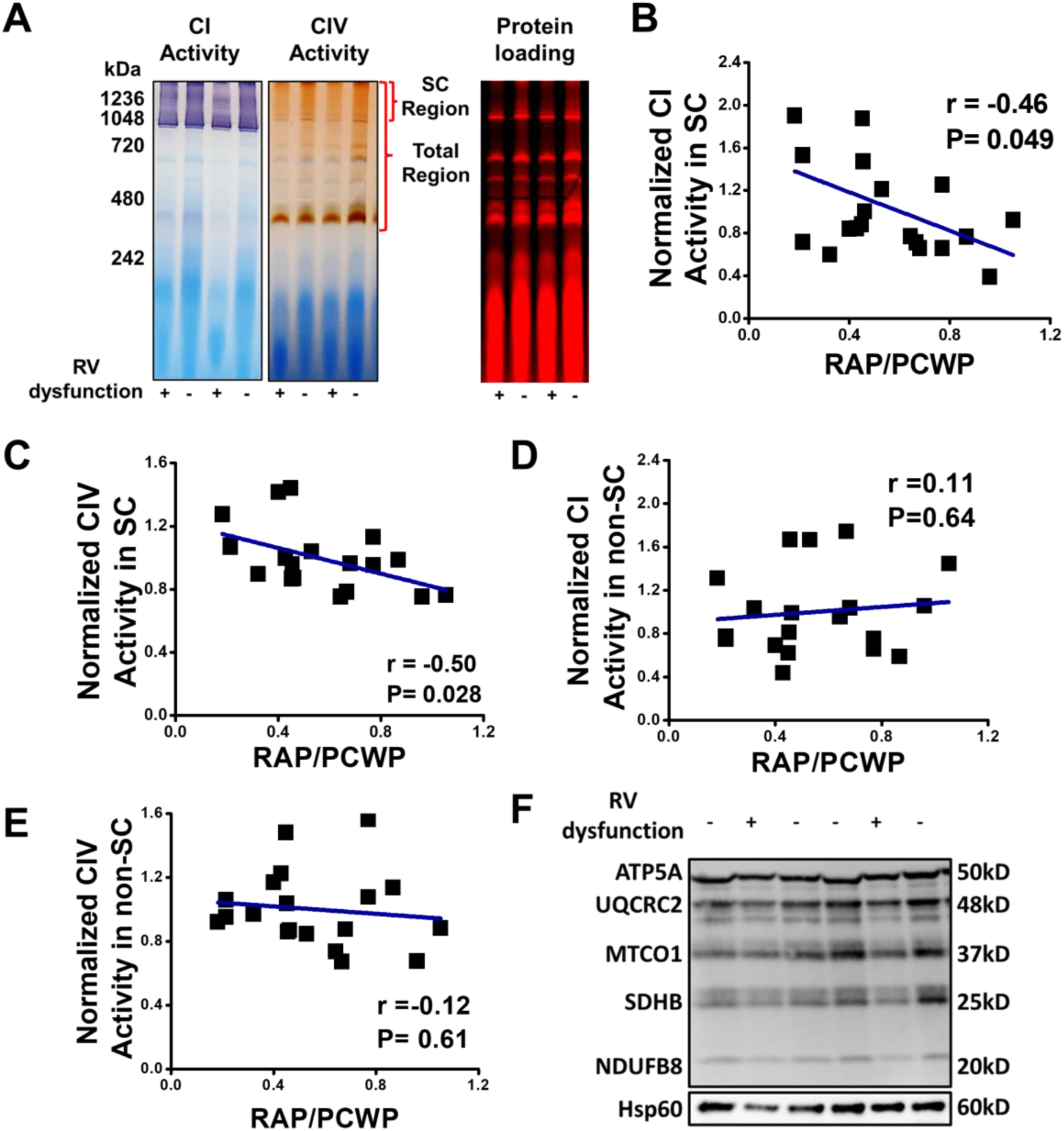
Explanted human hearts with RV dysfunction have reduced mSC activity of CI and CIV in the RV. **A**) Representative CN-PAGE of CI and CIV activity in human RV from explanted hearts, and licor far red imaging of Coomassie stain for protein load. + indicates high RAP/PCWP (>.7) and – low RAP/PCWP (<.3). **B,C**) CI and CIV activity in mSC region negatively correlates with RAP/PCWP in human RV (high RAP/PCWP indicates RV dysfunction). **D,E**) CI and CIV activity in the non-mSC region does not significantly correlate with RAP/PCWP in human RV. **F**) Representative SDS-PAGE blot of individual ETC complex subunits from RV tissue. There was no significant correlation with RAP/PCWP (not shown). R and P values determined by Pearson’s correlation, n=19.

**Table 2:** Patient demographics and RV functional parameters of explanted human heart samples. Data presented as mean +/- S.D.

|  |  |
| --- | --- |
| **Age(median)** | 37(19-67) |
| **Sex (M,F)** | 8,11 |
| **Tobacco** | Current smoking (1),Smoked(9), Never smoked(7),Unknown (2) |
| **Heart failure type** | Non-ischemic cardiomyopathy(12), Peripartum cardiomyopathy(3), HFrEF(3) Cardiomyopathy(1) |
| **mPAP** | 30.00 ± 9.99 |
| **PCWP** | 20.26 ± 9.24 |
| **RAP** | 10.79 ± 6.06 |
| **Cardiac Index** | 2.18 ± 0.61 |
| **RVSWI** | 7.67 ± 3.59 |
| **RAP/PWCP** | 0.55 ± 0.25 |

mPAP: mean pulmonary arterial pressure, PCWP: pulmonary capillary wedge pressure, RAP: right atrial pressure, RVSWI: right ventricular stroke work index

## Discussion

This study is the first to demonstrate pronounced deficits in mitochondrial mSC activity and assembly associated with RV dysfunction in both the sugen/hypoxia rat model of PH as well as in humans with RV dysfunction due to group II PH. Bioenergetic and metabolic changes during RV hypertrophy and failure in response to PH are increasingly recognized. RV dysfunction is associated with significant alterations to mitochondrial function and bioenergetics. Mitochondrial changes that have been described in the RV in PH include altered mitochondrial metabolic gene expression, decreased respiration, mitochondrial structural and cristae remodeling, and oxidant stress all of which may impair RV function[8, 18, 40, 41] . The RV has been shown to undergo metabolic reprogramming with a decrease in oxidative phosphorylation and increase in glycolysis and glutaminolysis likely to sustain energy production[18, 28]. Here we demonstrate that, in addition to reductions in mitochondrial content, ETC protein expression, or changes in energy utilization pathways, there are significant reductions in mitochondrial SC assembly and a marked reduction in mSC activity in the RV in PH. These changes in mSCs will need to be considered as a potentially important contributor to mitochondrial dysfunction among the already recognized pathologic bioenergetic changes in PH, as reductions in mSCs will likely contribute to decreased ATP production and promote more inefficient respiration and increased oxidative stress.

It is of note that only changes in activity of mSC’s correlate with impaired RV function in PH rats and humans with RV dysfunction, and not total CI and CIV activity associated with individual or smaller non-supercomplex subassemblies (such as I/III and III/IV assemblies) as shown in **Table 1** and **Figure 7**. There is considerable evidence to support the relative importance of mSC-mediated respiration in the “solid state” or electron tunneling models (supercomplex only mediated respiration) versus more dynamic cellular respiration utilizing non-SC assemblies and intermembrane diffusion of complex subunits for electron transfer [1, 3, 10, 15, 19, 21]. Our data suggests that in the energetically demanding RV cardiac muscle, reductions in mSC activity and assembly may be of critical importance causing impaired respiration that may contribute to RV dysfunction and failure.

Loss of mSC-mediated respiration may promote more inefficient respiration through individual ETC complexes to meet energy demand at the cost of increased ROS [24]. This may lead to subsequent cellular damage leading to RV dysfunction and failure. Indeed, we show PH rat RV myocytes exhibited a greatly enhanced mitochondrial ROS generation as assessed by matrix-targeted HyPer7 ratios as well as reductions in mitochondrial membrane potential (**Figure 6**), despite RV myocardium having relatively normal expression of subunits of the ETC and activity and assembly of smaller ETC complex assemblies (**Figures 2-5**). Despite numerous studies that have identified PH-induced changes in RV redox sensitive/ antioxidant proteins or cascades suggesting elevated ROS[13, 35, 41], there have been extremely limited studies that directly measured ROS generation in the RV myocardium or myocytes. Also, the prior studies have shown conflicting results, have only been performed in the MCT model and none have directly measured RV cardiomyocyte mitochondrial-specific ROS. Redout et al. found greatly increased superoxide using spectrophotometric methods of RV homogenates and isolated mitochondria from the monocrotaline rat model, Zimmer et al. showed mild increases at 1 week that resolve in the MCT model using spectrophotometric methods on RV tissue lysates and, Puukila et al, showed no changes in tissue lysates measured with the general cytoplasmic dye DCF-DA [30, 31, 43]. Here, we demonstrate a robust increase (∼50% baseline increases) in mitochondrial ROS generation in RV cardiomyocytes in the SuHx model that parallels the decrease in mSC. Our experiments using matrix-HyPer7 have a number of advantages over methods employed by others previously including 1) imaging in live cells, 2) the highly sensitive new generation ROS sensor is genetically targeted to mitochondria, 3) observations are ratiometric thereby nullifying any dye loading or variable expression concerns, and 4) the ratios are quantifiable following normalization to minimal and maximal signals.

While the importance of mSC in mitochondrial function is well established, there have been limited studies on mSC derangements in heart failure in either the LV or the RV. Hoppel et al. demonstrated reductions in LV mSC in a dog intracoronary embolization HF model [32] and a recent study with explanted dilated cardiomyopathy hearts from young patients showed a non-significant reduction in mSC activity using in-gel activity assay [4]. There are multiple studies with conflicting results regarding the importance of mSC disassembly in multiple animal and isolated heart models of LV ischemic injury, with increased, decreased or unaltered mSC with ischemic injury [15]. However, it should be noted these studies generally interrogate acute ischemic injury with short time courses of injury in contrast to the human data presented here or our 7 week SuHx data. Javadov et. al, recently published an excellent review on I/R mSC studies and propose that the differing severity of I/R injury and significant variation in techniques used to assess mSC account for the conflicting results [15]. Studies of mSC in long term heart failure studies are much more limited. Genetic models of Barth Syndrome involving Tafazzin KO or KD have demonstrated there are decreases in mSC (due to cardiolipin disturbances) that associate with development of cardiomyopathy [9, 42]. However, there are gross mitochondrial structural alterations with Tafazzin KO/KD confounding clear interpretation regarding the importance of mSC assembly. In, addition knock down of NDUFAB1, a component of complex I that promotes CI and subsequently mSC assembly, results in impaired LV CI and mSC assembly, respiration and HF[12]. Overexpression of NDUFAB1 was protective for ischemic injury, however a protective role in HF is unknown. In the RV, Wust et al. showed reduced total activity of CI (inclusive of mSC and smaller CI assemblies) in a rat monocrotaline model of right heart failure (RHF), however, they did not find differences in CI incorporation into supercomplexes [40]. This is in contrast to our data in the longer duration sugen-hypoxia rat model of pulmonary arterial hypertension (PAH) and in humans with RV dysfunction, and may reflect differences in both the model and techniques employed by both studies.

Surprisingly, we identified significant decrease in total CIII assembly, reduced inclusion of CIII and CIV in mSC assemblies, and reduced activity of CIV in mSC in the LV of PAH rats (Fig 2 and 4). Although these changes appeared mild compared to the RV. There are numerous changes to the LV in PAH including altered chamber geometry, reduced filling, contraction, and LV atrophy [11, 23, 33] . It is possible that the reduced filling and atrophic signaling may play a role in altered mSC assembly and bioenergetic remodeling in the LV. We have previously found that ETC complexes were reduced in specific skeletal muscles in sedentary SuHx rats and ETC assembly could be restored with exercise [25]. Overall, our data suggests that alterations to mSC organization and activity may also play a role in changes to the LV in PAH, however the cause of these changes whether pathological or adaptive due to decreased cardiac output driven by RV dysfunction are unclear. We speculate that altered LV mSC may be associated with reduced metabolic demand due to reduced LV filling and cardiac output.

In conclusion, our data provide evidence that ETC SC assembly and activity are greatly reduced in the RV following chronic pressure overload in PH and in humans with RV dysfunction associated with pulmonary hypertension. We speculate that modulating mSC assembly and activity may be a viable therapeutic strategy in PH. Strategies to preserve or increase mSC may mitigate bioenergetic abnormalities and reduce ROS in the pressure overloaded RV to delay progression to heart failure. Future studies and further investigation will need to identify robust interventions that can safely boost mSC activity to determine if either driving mSC activity or limiting mSC derangements in the RV is beneficial in PH.

### Limitations

There are several limitations with the current manuscript that should be taken into consideration. First, ROS data is from separate animals as there are limited methods to detect ROS-dependent protein modification in tissue and which do not approach the sensitivity or specificity of ROS imaging in live myocytes. However, this does preclude correlational analysis with the detected derangements in RV supercomplex activity and assembly. Second, we did not have access to any RV samples from Group I PAH patients to compare with the SuHx rats a model of PAH, as heart/lung transplant in this patient population is exceptionally rare in most transplant centers compared to the majority of transplants associated with left heart disease. However, commonality of reduced SCs in both humans with RV dysfunction and the rat PAH model, would argue that this is common in RV maladaptation and dysfunction regardless of PH group classification.

## Abbreviations

PH: Pulmonary hypertension
mSC: mitochondrial supercomplex
ETC: electron transport chain
mPAP: mean pulmonary arterial pressure
RVH: right ventricular hypertrophy
ROS: reactive oxygen species
BN: Blue native
CN: clear native
TAPSE: tricuspid annular plane systolic excursion
PAT: pulmonary artery acceleration time
CO: cardiac output
TV E/E’: tricuspid valve peak velocity of early diastolic transmitral flow/ peak velocity of early diastolic mitral annular motion
PCWP: pulmonary capillary wedge pressure
RAP: right atrial pressure
RVSWI: right ventricular stroke work index
DTT: Dithiothreitol
TMRE: Tetramethylrhodamine ethyl ester.

## References

1. Acin-Perez R, Enriquez J a (2014) The function of the respiratory supercomplexes: The plasticity model. Biochimica et Biophysica Acta (BBA) - Bioenergetics 1837:444–450. doi: 10.1016/j.bbabio.2013.12.009

2. Blair CA, Haynes P, Campbell SG, Chung C, Mitov MI, Dennis D, Bonnell MR, Hoopes CW, Guglin M, Campbell KS (2016) A Protocol for Collecting Human Cardiac Tissue for Research. The VAD Journal 2. doi: 10.14434/vad.v2i0.27941

3. Calvo E, Cogliati S, Hernansanz-Agustín P, Loureiro-López M, Guarás A, Casuso RA, García-Marqués F, Acín-Pérez R, Martí-Mateos Y, Silla-Castro JC, Carro-Alvarellos M, Huertas JR, Vázquez J, Enríquez JA (2020) Functional role of respiratory supercomplexes in mice: SCAF1 relevance and segmentation of the Qpool. Sci Adv 6. doi: 10.1126/SCIADV.ABA7509

4. Chatfield KC, Sparagna GC, Chau S, Phillips EK, Ambardekar A V., Aftab M, Mitchell MB, Sucharov CC, Miyamoto SD, Stauffer BL (2019) Elamipretide Improves Mitochondrial Function in the Failing Human Heart. JACC Basic Transl Sci 4:147–157. doi: 10.1016/j.jacbts.2018.12.005

5. Clements RT, Terentyeva R, Hamilton S, Janssen PML, Roder K, Martin BY, Perger F, Schneider T, Nichtova Z, Das AS, Veress R, Lee BS, Kim DG, Koren G, Stratton MS, Csordas G, Accornero F, Belevych AE, Gyorke S, Terentyev D (2023) Sexual dimorphism in bidirectional SR-mitochondria crosstalk in ventricular cardiomyocytes. Basic Res Cardiol 118. doi: 10.1007/S00395-023-00988-1

6. Clements RT, Vang A, Fernandez-Nicolas A, Kue NR, Mancini TJ, Morrison AR, Mallem K, McCullough DJ, Choudhary G (2019) Treatment of Pulmonary Hypertension with Angiotensin II Receptor Blocker and Neprilysin Inhibitor Sacubitril/Valsartan. Circ Heart Fail 12. doi: 10.1161/CIRCHEARTFAILURE.119.005819

7. Drazner MH, Velez-Martinez M, Ayers CR, Reimold SC, Thibodeau JT, Mishkin JD, Mammen PPA, Markham DW, Patel CB (2013) Relationship of right- to left-sided ventricular filling pressures in advanced heart failure insights from the escape trial. Circ Heart Fail 6:264–270. doi: 10.1161/CIRCHEARTFAILURE.112.000204

8. Gomez-Arroyo J, Mizuno S, Szczepanek K, Van Tassell B, Natarajan R, Dos Remedios CG, Drake JI, Farkas L, Kraskauskas D, Wijesinghe DS, Chalfant CE, Bigbee J, Abbate A, Lesnefsky EJ, Bogaard HJ, Voelkel NF (2013) Metabolic gene remodeling and mitochondrial dysfunction in failing right ventricular hypertrophy secondary to pulmonary arterial hypertension. Circ Heart Fail 6:136–144. doi: 10.1161/CIRCHEARTFAILURE.111.966127/-/DC1

9. Greenwell AA, Tabatabaei Dakhili SA, Ussher JR (2022) Myocardial disturbances of intermediary metabolism in Barth syndrome. Front Cardiovasc Med 9. doi: 10.3389/FCVM.2022.981972

10. Gu J, Wu M, Guo R, Yan K, Lei J, Gao N, Yang M (2016) The architecture of the mammalian respirasome. Nature 537:639–643. doi: 10.1038/NATURE19359

11. Hardziyenka M, Campian ME, Reesink HJ, Surie S, Bouma BJ, Groenink M, Klemens CA, Beekman L, Remme CA, Bresser P, Tan HL (2011) Right ventricular failure following chronic pressure overload is associated with reduction in left ventricular mass: evidence for atrophic remodeling. J Am Coll Cardiol 57:921–928. doi: 10.1016/J.JACC.2010.08.648

12. Hou T, Zhang R, Jian C, Ding W, Wang Y, Ling S, Ma Q, Hu X, Cheng H, Wang X (2019) NDUFAB1 confers cardio-protection by enhancing mitochondrial bioenergetics through coordination of respiratory complex and supercomplex assembly. Cell Res 29:754–766. doi: 10.1038/S41422-019-0208-X

13. Hwang H V., Sandeep N, Nair R V., Hu DQ, Zhao M, Lan IS, Fajardo G, Matkovich SJ, Bernstein D, Reddy S (2021) Transcriptomic and functional analyses of mitochondrial dysfunction in pressure overload-induced right ventricular failure. J Am Heart Assoc 10:1–47. doi: 10.1161/JAHA.120.017835

14. Jani V, Aslam MI, Fenwick AJ, Ma W, Gong H, Milburn G, Nissen D, Cubero Salazar IM, Hanselman O, Mukherjee M, Halushka MK, Margulies KB, Campbell KS, Irving TC, Kass DA, Hsu S (2023) Right Ventricular Sarcomere Contractile Depression and the Role of Thick Filament Activation in Human Heart Failure With Pulmonary Hypertension. Circulation 147:1919–1932. doi: 10.1161/CIRCULATIONAHA.123.064717

15. Javadov S, Jang S, Chapa-Dubocq XR, Khuchua Z, Camara AK (2021) Mitochondrial respiratory supercomplexes in mammalian cells: structural versus functional role. J Mol Med (Berl) 99:57–73. doi: 10.1007/S00109-020-02004-8

16. Jha P, Wang X, Auwerx J (2016) Analysis of Mitochondrial Respiratory Chain Supercomplexes Using Blue Native Polyacrylamide Gel Electrophoresis (BN-PAGE). Curr Protoc Mouse Biol 6:1–14. doi: 10.1002/9780470942390.mo150182

17. Kazmirczak F, Hartweck LM, Vogel NT, Mendelson JB, Park AK, Raveendran RM, O- Uchi J, Jhun BS, Prisco SZ, Prins KW (2023) Intermittent Fasting Activates AMP-Kinase to Restructure Right Ventricular Lipid Metabolism and Microtubules. JACC Basic Transl Sci 8:239–254. doi: 10.1016/J.JACBTS.2022.12.001

18. Koop AMC, Bossers GPL, Ploegstra MJ, Hagdorn QAJ, Berger RMF, Silljé HHW, Bartelds B (2019) Metabolic Remodeling in the Pressure-Loaded Right Ventricle: Shifts in Glucose and Fatty Acid Metabolism—A Systematic Review and Meta-Analysis. J Am Heart Assoc 8

19. Lapuente-Brun E, Moreno-Loshuertos R, Acín-Pérez R, Latorre-Pellicer A, Colás C, Balsa E, Perales-Clemente E, Quirós PM, Calvo E, Rodríguez-Hernández M a, Navas P, Cruz R, Carracedo Á, López-Otín C, Pérez-Martos A, Fernández-Silva P, Fernández-Vizarra E, Enríquez JA (2013) Supercomplex Assembly Determines Electron Flux in the Mitochondrial Electron Transport Chain. Science (1979) 340:1567–1570. doi: 10.1126/science.1230381

20. Letts JA, Fiedorczuk K, Sazanov LA (2016) The architecture of respiratory supercomplexes. Nature 537:644–648. doi: 10.1038/nature19774

21. Letts JA, Sazanov LA (2017) Clarifying the supercomplex: The higher-order organization of the mitochondrial electron transport chain. Nat Struct Mol Biol 24:800–808. doi: 10.1038/NSMB.3460

22. Li X, Braza J, Mende U, Choudhary G, Zhang P (2021) Cardioprotective effects of early intervention with sacubitril/valsartan on pressure overloaded rat hearts. Sci Rep 11. doi: 10.1038/S41598-021-95988-3

23. Manders E, Bogaard HJ, Handoko ML, Van De Veerdonk MC, Keogh A, Westerhof N, Stienen GJM, Dos Remedios CG, Humbert M, Dorfmüller P, Fadel E, Guignabert C, Van Der Velden J, Vonk-Noordegraaf A, De Man FS, Ottenheijm CAC (2014) Contractile Dysfunction of Left Ventricular Cardiomyocytes in Patients With Pulmonary Arterial Hypertension. J Am Coll Cardiol 64:28–37. doi: 10.1016/J.JACC.2014.04.031

24. Maranzana E, Barbero G, Falasca AI, Lenaz G, Genova ML (2013) Mitochondrial Respiratory Supercomplex Association Limits Production of Reactive Oxygen Species from Complex I. Antioxid Redox Signal 19:1469–1480. doi: 10.1089/ars.2012.4845

25. McCullough DJ, Kue N, Mancini T, Vang A, Clements RT, Choudhary G (2020) Endurance exercise training in pulmonary hypertension increases skeletal muscle electron transport chain supercomplex assembly. Pulm Circ 10. doi: 10.1177/2045894020925762

26. Pak V V., Ezeriņa D, Lyublinskaya OG, Pedre B, Tyurin-Kuzmin PA, Mishina NM, Thauvin M, Young D, Wahni K, Martínez Gache SA, Demidovich AD, Ermakova YG, Maslova YD, Shokhina AG, Eroglu E, Bilan DS, Bogeski I, Michel T, Vriz S, Messens J, Belousov V V. (2020) Ultrasensitive Genetically Encoded Indicator for Hydrogen Peroxide Identifies Roles for the Oxidant in Cell Migration and Mitochondrial Function. Cell Metab 31:642–653.e6. doi: 10.1016/J.CMET.2020.02.003

27. Piao L, Fang Y-H, Cadete VJJ, Wietholt C, Urboniene D, Toth PT, Marsboom G, Zhang HJ, Haber I, Rehman J, Lopaschuk GD, Archer SL The inhibition of pyruvate dehydrogenase kinase improves impaired cardiac function and electrical remodeling in two models of right ventricular hypertrophy: resuscitating the hibernating right ventricle. doi: 10.1007/s00109-009-0524-6

28. Piao L, Fang YH, Parikh K, Ryan JJ, Toth PT, Archer SL (2013) Cardiac glutaminolysis: A maladaptive cancer metabolism pathway in the right ventricle in pulmonary hypertension. J Mol Med 91:1185–1197. doi: 10.1007/s00109-013-1064-7

29. Prisco SZ, Thenappan T, Prins KW (2020) Treatment Targets for Right Ventricular Dysfunction in Pulmonary Arterial Hypertension. JACC Basic Transl Sci 5:1244. doi: 10.1016/J.JACBTS.2020.07.011

30. Puukila S, Rafael ·, Fernandes O, Türck P, Campos Carraro C, Jéssica ·, Poletto Bonetto H, Gazzi De Lima-Seolin B, Sander A, Araujo R, Belló-Klein A, Boreham D, Khaper N (2017) Secoisolariciresinol diglucoside attenuates cardiac hypertrophy and oxidative stress in monocrotaline-induced right heart dysfunction. Mol Cell Biochem 432:33–39. doi: 10.1007/s11010-017-2995-z

31. Redout EM, Wagner MJ, Zuidwijk MJ, Boer C, Musters RJP, van Hardeveld C, Paulus WJ, Simonides WS (2007) Right-ventricular failure is associated with increased mitochondrial complex II activity and production of reactive oxygen species. Cardiovasc Res 75:770–781. doi: 10.1016/J.CARDIORES.2007.05.012

32. Rosca MG, Vazquez EJ, Kerner J, Parland W, Chandler MP, Stanley W, Sabbah HN, Hoppel CL (2008) Cardiac mitochondria in heart failure: decrease in respirasomes and oxidative phosphorylation. Cardiovasc Res 80:30–9. doi: 10.1093/cvr/cvn184

33. Rosenkranz S, Howard LS, Gomberg-Maitland M, Hoeper MM (2020) Systemic Consequences of Pulmonary Hypertension and Right-Sided Heart Failure. Circulation 141:678–693. doi: 10.1161/CIRCULATIONAHA.116.022362

34. Ryan JJ, Archer SL (2014) The right ventricle in pulmonary arterial hypertension: Disorders of metabolism, angiogenesis and adrenergic signaling in right ventricular failure. Circ Res 115:176–188. doi: 10.1161/CIRCRESAHA.113.301129

35. Schlüter KD, Kutsche HS, Hirschhäuser C, Schreckenberg R, Schulz R (2018) Review on chamber-specific differences in right and left heart reactive oxygen species handling. Front Physiol 9

36. Sikkel MB, Francis DP, Howard J, Gordon F, Rowlands C, Peters NS, Lyon AR, Harding SE, Macleod KT (2017) Hierarchical statistical techniques are necessary to draw reliable conclusions from analysis of isolated cardiomyocyte studies. Cardiovasc Res 113:1743– 1752. doi: 10.1093/CVR/CVX151

37. Taraseviciene-Stewart L, Kasahara Y, Alger L, Hirth P, Mc Mahon G, Waltenberger J, Voelkel NF, Tuder RM (2001) Inhibition of the VEGF receptor 2 combined with chronic hypoxia causes cell death-dependent pulmonary endothelial cell proliferation and severe pulmonary hypertension. FASEB J 15:427–38. doi: 10.1096/fj.00-0343com

38. Vang A, Clements RT, Chichger H, Kue N, Allawzi A, O’Connell K, Jeong EM, Dudley SC, Sakhatskyy P, Lu Q, Zhang P, Rounds S, Choudhary G (2017) Effect of α7 nicotinic acetylcholine receptor activation on cardiac fibroblasts:A mechanism underlying RV fibrosis associated with cigarette smoke exposure. Am J Physiol Lung Cell Mol Physiol 312:L669–L677. doi: 10.1152/ajplung.00393.2016

39. van de Veerdonk MC, Bogaard HJ, Voelkel NF (2016) The right ventricle and pulmonary hypertension. Heart Fail Rev. doi: 10.1007/s10741-016-9526-y

40. Wüst RCI, de Vries HJ, Wintjes LT, Rodenburg RJ, Niessen HWM, Stienen GJM (2016) Mitochondrial complex I dysfunction and altered NAD(P)H kinetics in rat myocardium in cardiac right ventricular hypertrophy and failure. Cardiovasc Res 111:362–72. doi: 10.1093/cvr/cvw176

41. Xu W, Janocha AJ, Erzurum SC (2021) Metabolism in Pulmonary Hypertension. Annu Rev Physiol 83:551. doi: 10.1146/ANNUREV-PHYSIOL-031620-123956

42. Zhu S, Chen Z, Zhu M, Shen Y, Leon LJ, Chi L, Spinozzi S, Tan C, Gu Y, Nguyen A, Zhou Y, Feng W, Vaz FM, Wang X, Gustafsson AB, Evans SM, Kunfu O, Fang X (2021) Cardiolipin Remodeling Defects Impair Mitochondrial Architecture and Function in a Murine Model of Barth Syndrome Cardiomyopathy. Circ Heart Fail 14:E008289. doi: 10.1161/CIRCHEARTFAILURE.121.008289

43. Zimmer A, Teixeira · R B, Bonetto · J H P, Bahr · A C, Türck · P, De Castro · A L, Campos-Carraro · C, Visioli · F, Fernandes-Piedras · T R, Casali · K R, Scassola · C M C, Baldo · G, Araujo · A S, Singal · P, Belló-Klein · A (2020) Role of inflammation, oxidative stress, and autonomic nervous system activation during the development of right and left cardiac remodeling in experimental pulmonary arterial hypertension. Mol Cell Biochem 464:93–109. doi: 10.1007/s11010-019-03652-2

